# A Unified Framework for Lineage Tracing and Trajectory Inference

**DOI:** 10.1101/2020.07.31.231621

**Authors:** Aden Forrow, Geoffrey Schiebinger

## Abstract

Understanding the genetic and epigenetic programs that control differentiation during development is a fundamental challenge, with broad impacts across biology and medicine. New measurement technologies like single-cell RNA-sequencing and CRISPR-based lineage tracing have opened new windows on these processes, through computational trajectory inference and lineage reconstruction. While these two mathematical problems are deeply related, they been approached from separate directions: methods for trajectory inference are not typically designed to leverage information from lineage tracing and vice versa. We present a novel, unified framework for lineage tracing and trajectory inference. Specifically, we develop a method for reconstructing developmental trajectories from time courses with snapshots of both cell states and lineages, leveraging mathematical tools from graphical models and optimal transport. We find that lineage data helps disentangle complex state transitions with increased accuracy using fewer measured time points. Moreover, integrating lineage tracing with trajectory inference in this way enables accurate reconstruction of developmental pathways that are impossible to recover with state-based methods alone.

## Introduction

Analyzing the trajectories of cellular differentiation holds promise for key questions across biology, from how lineages diverge during embryonic development to how cell types destabilize with age or are perturbed in disease. Single-cell measurement technologies like single-cell RNA-sequencing (scRNA-seq) [9, 13], single-cell ATAC-seq [2], and CRISPR-based lineage tracing [14, 19, 22] have opened new windows on these processes, but it remains challenging to analyze dynamic changes in cell state and cell lineage over time because the measurements are destructive: cells must be lysed before information about their state or lineage can be recovered. In response, there has been a flurry of work on designing methods to infer developmental trajectories from static snapshots of cell state [5, 28, 35, 36], including our own efforts [24]. While initial efforts have shed some light on important biological questions relating to embryonic development [1, 5], hematopoiesis [34], and induced pluripotent stem cell reprogramming [24], the field of trajectory inference is still in its infancy.

One of the most significant deficiencies of existing trajectory inference methods is that they are not designed to incorporate the rich information from lineage-tracing. Technologies for reconstructing cellular lineage trees have seen tremendous recent advances, fueled by the CRISPR–Cas9 genome editing technology [4, 19, 22]. While developmental biologists have long used various methods to tag cells and trace the lineage of their descendants, newer approaches make it possible to recover more complex lineage relationships, including the full lineage tree of a population of cells [14, 19, 22]. These newer technologies employ CRISPR-Cas9 to continuously mutate an array of synthetic DNA barcodes, which are incorporated into the chromosomes so that they are inherited by daughter cells and can be further mutated over the course of development. By analyzing the pattern of mutations in the barcodes, one can reconstruct a lineage tree describing shared ancestry within a population of cells. Recent advances allow the DNA barcodes to be expressed as transcripts and recovered together with the rest of the transcriptome in scRNA-seq [19, 22]. Alternative methods use somatic mutations in mitochondria [12] to recover similar information without needing to introduce DNA barcodes.

Each of these technologies enables simultaneous collection of data on cell state and cell lineage, which provides an experimental solution to part of the trajectory inference problem. High-resolution lineage tracing, however, does not obviate the need for trajectory inference because lineage tracing alone does not reveal the state of the ancestral cells. While the problems of reconstructing lineage trees and inferring trajectories have attracted substantial attention individually [20, 33], there is much to be gained from combining these two complementary perspectives [6].

Here, we propose an integrated mathematical framework for inferring developmental trajectories from snapshots of both cell lineage and cell state. Our framework, called LineageOT, is broadly applicable to lineage tracing time-courses, where populations of cells are profiled with both scRNA-seq and lineage tracing at various time points along a developmental process (Fig. 1). As a proof of concept, we test our methodology on a time-course of *C. elegans* embryonic development (Figs. 2-3), collected with scRNA-seq [17]. Because the lineage tree of *C. elegans* is known [29], we have an objective measure of performance. We find that our method significantly improves trajectory inference both on this data set and on simulated examples where algorithms without lineage information cannot completely recover the correct trajectories (Fig. 4). Our results show a path towards realizing the substantial benefits of lineage tracing [6, 32] in applications across developmental biology.

**Figure 1:**
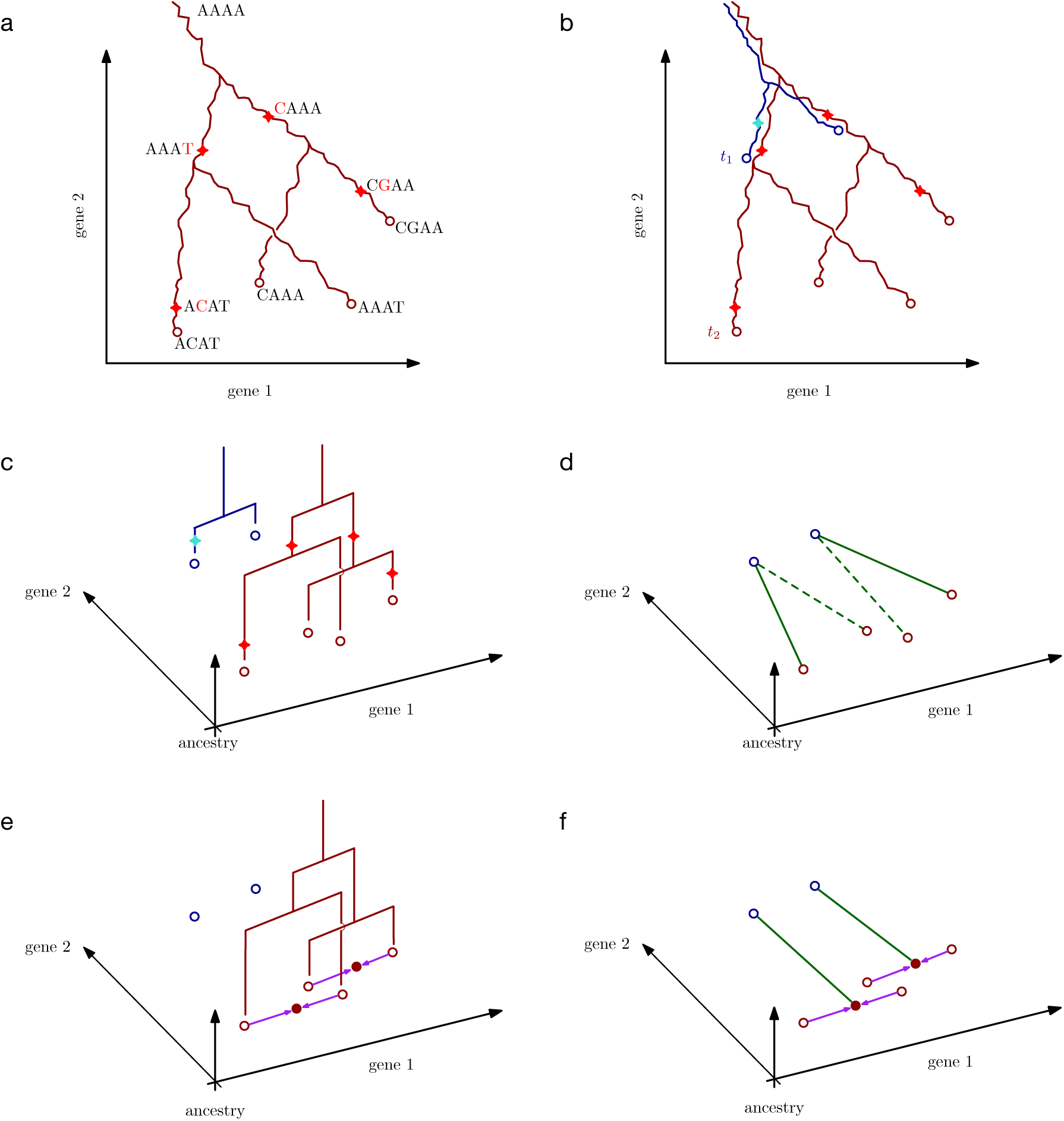
Schematic of the LineageOT model and inference procedure. (a) A lineage tree embedded in two dimensional gene expression space. As cells change state over time, they trace out paths. Branches in the tree correspond to cell divisions, giving rise to four cells at the measurement time (red circles). Each cell has a barcode to track its lineage. Starting from the ancestral barcode sequence *AAAA*, mutations are indicated with a red star on the lineage tree and the change to the sequence is shown in red. (b) Embedded lineage trees from two independent realizations of the developmental process measured at times *t*_1_ (blue) and *t*_2_ (red). (c) The setup from (b) is shown in a 3d plot with lineage trees visualized in the vertical dimension. For each time point, we observe cell states (dots) and also the lineage tree, but not the lineage tree embedded in state space. (d) A purely state-based algorithm would fail to recover the correct trajectories in this example. Green lines connect cells at *t*_2_ to their nearest neighbor at *t*_1_. Dashed lines indicate erroneous connections. (e-f) The LineageOT procedure consists of two steps. (e) Adjust cells at time *t*_2_ (purple arrows), based on lineage information. Cells with shared lineage are moved closer together, towards an estimate of ancestral state (solid dots). (f) Infer a coupling (green lines) connecting the adjusted cells from time *t*_2_ (red) to cells from time *t*_1_ (blue). This corrects the mistake made in (d).

**Figure 2:**
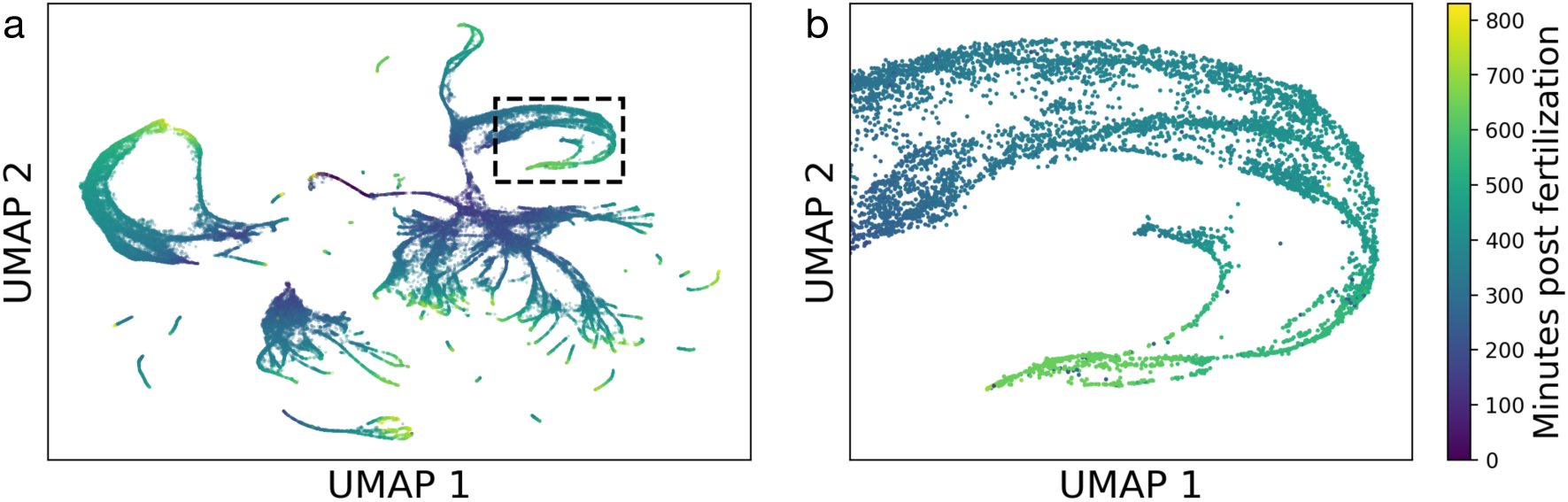
(a) UMAP of 81286 *C. elegans* cells from [17], using coordinates provided by Packer et al. Color indicates estimated time since fertilization following the colorbar in (b). (b) In the boxed region from (a), multiple developmental trajectories in the hypodermis converge to the same UMAP coordinates, suggesting a convergence in gene expression.

**Figure 4:**
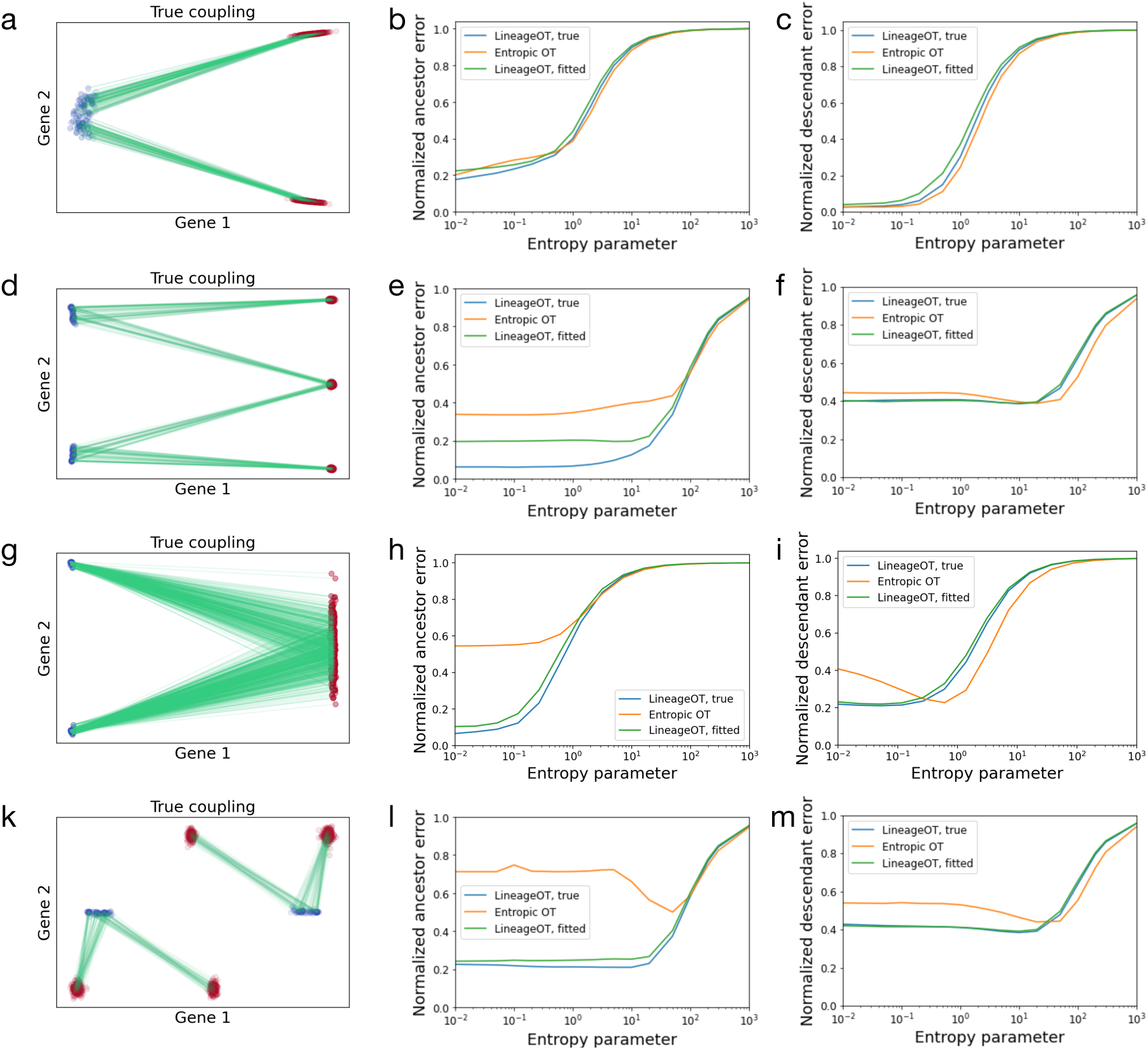
LineageOT matches the performance of Waddington-OT for simple trajectories and exceeds it for complex trajectories. (a-c) For a simple bifurcation, optimal transport alone works well and adding lineage information makes little difference. (a) We simulated a cluster of cells at an early time point splitting into two clusters at a later time point. Green lines connect ancestors in blue to descendants in red in (a), (d), (g) and (k). The ancestor errors (b) and descendant errors (c) are similar for optimal transport (orange) and LineageOT (blue) with any entropy parameter, even when LineageOT is given an imperfect tree fitted to simulated barcodes (green). (d-f) For a convergent trajectory, LineageOT significantly improves ancestor prediction with no loss of accuracy in descendant prediction, even with an imperfectly fitted lineage tree. (d) Here we simulated two early clusters that each split; later, two of the resulting clusters merge together. Using LineageOT reduces error substantially for ancestor prediction (e) and slightly for descendant prediction (f). (g-i) The improvement due to lineage information when trajectories converge does not require nearby unconverged clusters. Here we see qualitatively similar improvement for two early clusters whose distributions of descendant cells almost entirely overlap. (k-m) With sufficient time between samples, clusters of cells may move closer to early time point cells that are not their ancestors. (k) In this simulation, after two early clusters each split, two of the late clusters are closer to non-ancestral cells than to their true ancestors. Optimal transport couples clusters incorrectly, leading to high error for predicting both ancestors (l) and descendants (m). LineageOT corrects the errors in this example by averaging with other clusters that are mapped correctly.

## Results

### A unified framework for lineage tracing and trajectory inference

We develop a mathematical framework for analyzing scRNA-seq time-courses equipped with a lineage tree at each time point. We formulate the goal of trajectory inference in terms of recovering the embedding of these lineage trees, defined as follows. As a population of cells develops, each cell traces out trajectories in a high-dimensional vector space of cellular states (e.g. gene expression space). Cell divisions create branching paths, and the trajectories of related cells coincide up to the point when their ancestry diverges (Fig. 1a). For example, if all the cells share a common ancestor, then the trajectories will all originate from a common point. This collection of branching paths forms the embedded lineage tree for the population. Note the emphasis on ‘embedded’ — without this modifier, the term ‘lineage tree’ refers to the coordinate-free tree structure, where all information about the embedded state of each ancestral node is lost, like those in Fig. 1c.

Single cell measurement technologies allow us to sample from a population and measure cell states together with barcodes that enable recovery of the lineage tree any point in time (Fig. 1a). However, because the measurements are destructive, we cannot directly chart the embedded lineage tree at multiple time points. One can, however, leverage the reproducibility of development and collect samples from separate populations at different time points (Fig. 1b,c). For example, one can prepare two independent populations of cells and collect samples from the first population at time *t*_1_ and samples from the second population at time *t*_2_. The key question is then: *which cell from the first sample would have given rise to each cell from the second sample, if these were two views of the same population*?

Importantly, this cannot be solved in general from the topology of the lineage trees alone. Both biological variability and simple subsampling of cells in a tissue could cause the lineage tree at *t*_2_ to be topologically distinct from an extension of the lineage tree at *t*_1_, making it impossible to directly patch one tree onto the other in a biologically meaningful way. We sketch a hypothetical example with sampling in Fig. S1. Instead, lineage information must be used together with gene expression data in a combined approach.

We have recently demonstrated [24] that a classical mathematical tool called optimal transport [8, 15, 31] can be applied to infer ‘state couplings’ from a scRNA-seq time-course, without any information about cell lineage. This method, called Waddington-OT, connects cells sampled at time *t*_1_ to their putative descendants at time *t*_2_ by minimizing the total distance traveled by all cells. It also includes entropic regularization with a tunable regularization parameter to model the inherent stochasticity in developmental trajectories and allows for variable rates of growth across cells by adjusting the distributions at times *t*_1_ and *t*_2_ based on estimates of division rates. The inferred connections approximate the frequency of transitions between regions of cell-state space, i.e. the couplings of the developmental process.

Our present notion of an embedded lineage tree refines the notion of a coupling from [24]. Where Waddington-OT aims for a state coupling describing all possible ancestries of a hypothetical cell with a given state, our embedded lineage tree gives rise to a lineage-resolved coupling. The difference is significant in situations where cells can arrive at a particular state from different ancestral states (Methods 1). Lineage tracing helps resolve such ambiguities: without lineage tracing, we must assume that cells with similar states have similar ancestral states; with lineage tracing, we instead assume that cells with similar *lineage* have similar ancestral states.

We apply optimal transport to recover lineage couplings, considered as approximations to embedded lineage trees, from scRNA-seq time-courses equipped with an unembedded lineage tree at each time point. We refer to these datasets as scRNA-lineage time-courses. In practice, these unembedded trees can be reconstructed from mutations accumulated in DNA barcodes over the course of development (Fig. 1a), or some lineage information might be known in advance (as in *C. elegans* development). We do not focus on how the unembedded lineage trees are estimated – our method assumes these are calculated separately, for example by neighbor-joining [21], and given as input to the algorithm. However, we do demonstrate in simulations below that our method is robust to errors in the estimated lineage tree.

Our method applies two key steps to recover the lineage coupling spanning a pair of time points *t*_1_, *t*_2_.

1. We first leverage the lineage tree to adjust the positions of the cells at time *t*_2_(Fig. 1e).
2. We then connect them to their ancestors at time *t*_1_ (Fig. 1f). using entropically regularized optimal transport.

The adjustment in the first step can be interpreted as sharing information between closely related cells in order to construct a rough initial estimate of the ancestral states at the earlier time *t*_1_. The rationale behind the second step is based on Schrödinger’s discovery that entropically regularized optimal transport gives the maximum likelihood coupling of diffusing particles [11, 25]. Assuming that the trajectories are generated by diffusion plus drift through Waddington’s landscape (Methods 2), our estimates of ancestral states from the first step are approximately normally distributed, as we explain below. Therefore our procedure gives an approximate maximum likelihood estimate of the lineage coupling.

We now describe these two steps in detail. The core problem involves a single pair of time points, *t*_1_ and *t*_2_, where we are given cells *x*_1_, …, *x*_*n*_ sampled at time *t*_1_, and cells *y*_1_, …, *y*_*m*_ sampled at time *t*_2_ together with an estimate of their lineage tree. Because diffusion dominates drift on short time-scales, we can infer the ancestral state of *y*_*i*_ at time *t*_1_ by assuming the dynamics are driven by pure diffusion. However, conditional on the lineage tree, the cells are not diffusing independently. Intuitively, cells with similar lineage should diffuse back towards one another to reach a common ancestral state. The difference in cell state across each of the edges of the lineage tree is given by an independent Gaussian random variable with variance proportional to the time-span along the edge (Methods 4). This implies that the ancestral state at time *t*_1_ for each *y*_*i*_ is normally distributed with mean and variance that can be calculated from the lineage tree (Methods 4). Because the ancestral states of each *y*_*i*_ are normally distributed, optimal transport will give the maximum likelihood matching to the observed ancestors *x*_1_, …, *x*_*n*_, when we use an entropy parameter proportional to the inferred variance of ancestral states (Methods 4). This matching, or lineage-resolved coupling, summarizes our knowledge of the ancestral states of cells from *t*_2_ and the hypothetical descendant states of cells from *t*_1_, providing a window onto the embedded lineage tree of each time point.

As described in the original Waddington-OT paper [24], couplings of the kind fit by LineageOT reveal trajectory information covering the entire period between the earliest and latest sampling times. In between measured time points, trajectories can be estimated with geodesic interpolation, while concatenating a sequence of pairwise couplings between consecutive time points gives connections across longer periods. Moreover, in cases where the lineage tree at *t*_2_ is a direct extension of the lineage tree at *t*_1_, an accurately recovered coupling provides precisely the matching required to patch the tree topologies. The foundation for all of these analyses is a good performance for two time points, which we demonstrate for LineageOT in the subsequent sections.

### LineageOT outperforms Waddington-OT on a lineage-resolved time-course of *C. elegans* embryonic development

We sought to test our method by applying it to a scRNA-lineage time-course. While CRISPR-based lineage tracing [3, 26] offers tremendous potential for generating scRNA-lineage time-courses, this type of dataset has not yet been published. We reasoned, however, that we could create a scRNA-lineage time-course from an ordinary, non-barcoded, scRNA-seq time-course of *C. elegans* embryonic development [17], because the lineage tree is entirely known [29]. Packer et al. sampled 86,024 cells with 10X from loosely synchronized embryos spanning the first 800 minutes of *C. elegans* embryonic development. As visible in a UMAP embedding (Fig. 2a), the differentiation process is complex. In addition to the many branchings, the gene expression of distinct transient cell types converges for several tissues, including the hypodermis (Fig. 2b) and IL1/IL2 neurons (Fig. 4a of [17]). Such convergences cause difficulties for state-based trajectory inference, because cells with similar measured state have different histories.

Because the precise timing of each embryo is not known, Packer et al. estimated the developmental time of each cell by correlating gene expression levels with data from a previous bulk RNA time-course [7, 17]. They then divided the cells into groups with similar estimated developmental times. We treat these groups of cells as discrete time points along a scRNA-seq time-course, using the end of each group’s time interval as the group’s time of sampling.

To obtain the scRNA-lineage time-course required for LineageOT, we needed to incorporate lineage information at each time point. However, lineage annotations are missing from 46% of the cells in the Packer et al. dataset. Moreover, the lineage of many of the annotated cells is not completely specified: some symmetric lineages are not distinguished (e.g. cells whose true lineage is ABprp or ABplp are all labeled as ABpxp). We explored three different strategies to get around this problem of incomplete lineage information. We first simply filtered out all cells with imperfect lineage annotation. This leaves us with only 5123 cells but with no ambiguity in the lineage tree. Second, we restricted attention to the well-annotated ABpxp sublineage, which contains 7087 cells (entirely distinct from the 5123 cells above), and we treated the lineages ABprp and ABplp as if they were identical. Third, we filtered out only cells completely lacking lineage annotation. For cells with incomplete annotations, we imputed a precise lineage label by randomly selecting from the options consistent with the partial annotation. For each approach, we also removed a small number of cells (< 5%) whose assigned sampling time was before their birth time according to the reference lineage tree. These three strategies yield three scRNA-lineage datasets which we analyze separately. The results we describe below are broadly similar for each of the three strategies (Fig. 3, Fig. S2, Fig. S3).

**Figure 3:**
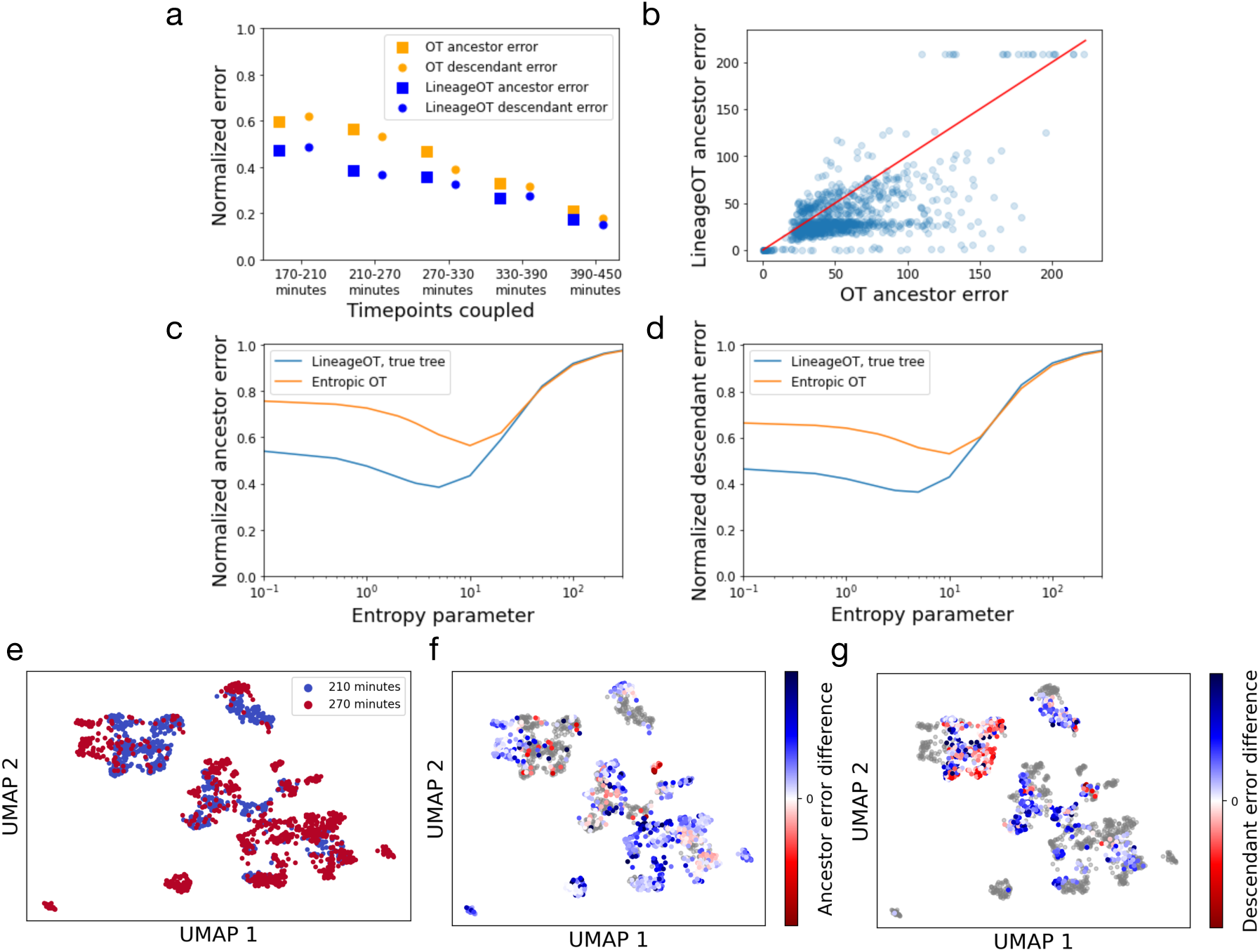
When tested on lineage-labeled *C. elegans* data, LineageOT outperforms optimal transport with no lineage information. (a) Relative accuracy of optimal transport and LineageOT on the 5123 cells with complete lineage annotations. Errors were normalized by dividing by the error of the noninformative independent coupling. (b) The error in predicting ancestor states, like the error for predicting descendant states (Fig. S4), is lower for most cells with LineageOT. Here each point represents one cell from the 270 minute time point, which was coupled to the 210 minute time point. The red line marks equal error for both methods. For each method in both (a-b) and (f-g), we chose the entropy parameters that gave the minimum error from parameter scans like those in (c) and (d). LineageOT consistently improves on Waddington-OT for reasonable values of the entropy parameter, both in ancestor error (c) and descendant error (d), shown here for the 210-270 minute couplings. (e) UMAP visualization of the cells from the 210 (blue) and 270 minute (red) time points. (f-g) Here, in the same UMAP, cells are colored by the ancestor (f) or descendant (g) error from Waddington-OT minus the same error from LineageOT. Blue indicates better performance by LineageOT, red better performance by Waddington-OT. The cells from 210 minutes and 270 minutes in (f) and (g) respectively are grey, as the corresponding error metric does not apply to them.

With each strategy, we applied both Waddington-OT [24] and LineageOT to infer developmental trajectories and compare their performance. We provide both methods with ground-truth growth rates (Methods 7), and compute state couplings and lineage couplings connecting each pair of time points. The input cell states are the first 50 coordinates from principal components analysis of normalized, log-transformed counts for the 46,159 cells with partial lineage annotations, corrected for background counts as in [17]. For LineageOT, the input lineage trees come from mapping cell lineage annotations onto the known *C. elegans* lineage tree. This additional information, given to LineageOT and not Waddington-OT, is precisely the information now measurable with lineage tracing.

We compare each fitted coupling to a ground-truth lineage coupling computed directly from the lineage-annotated data. This ground truth is constructed by connecting each early cell to all late cells labeled as being its descendants. Creating a coupling in this way would not be possible in other organisms without cell annotations from a known, invariant lineage tree. While previous work [20] has measured the success of trajectory inference by reducing to discrete branching representations, we directly check whether the predicted ancestors and descendants are similar in state to the true ancestors and descendants, respectively (Methods 6). These are two separate error metrics: the ancestor prediction error and the descendant prediction error. As an alternative, performance can be evaluated by the probability ancestor-descendant pairs from the ground truth are correctly linked. Though LineageOT does improve on Waddington-OT by that metric (Fig. S5), we prefer the ancestor and descendant error metrics because they do not require assuming the lineage trees from the two time points match.

In all our tests, LineageOT has consistently lower error for both ancestor and descendant prediction at reasonable levels of entropy (Fig. 3a, Fig. S2, Fig. S3), including after concatenating couplings across more than two time points (Fig. S6). LineageOT systematically predicts better for the majority of cells (Fig. 3b). The degree of improvement depends on the choice of entropic regularization parameter and the strategy for getting complete lineage annotations (Fig. 3c-d, Fig. S2, Fig. S3), but there is no entropy choice for which LineageOT performs significantly worse. The increased accuracy comes from effectively using the information in the lineage tree.

### Lineage-tracing trajectory inference outperforms state-based trajectory inference on complex trajectories

We next explored the performance of LineageOT on simulated data, with the goal of characterizing some of the settings where lineage-based trajectory inference can significantly outperform state-based trajectory inference. We found that lineage information is most helpful in resolving *convergent* trajectories, where similar cells arise from different ancestral states. Moreover, we found that LineageOT is robust to imperfections in the lineage tree. Below we present four simulations illustrating these concepts.

In each simulation, we generate an embedded lineage tree by allowing an initial population of cells to follow a vector field with diffusion and also to divide (Methods 8). Each cell has a lineage barcode that randomly mutates and is inherited by the cell’s descendants. We sample populations of cells at two time points, compute couplings with Waddington-OT and LineageOT, and compare to the ground truth coupling from the simulation, using the ancestor and descendant prediction errors we described above. We also test the robustness of LineageOT by giving the algorithm either (a) a lineage tree constructed from the simulated barcodes using a heuristic algorithm called neighbor-joining [21] (Methods 5) or (b) the ground-truth lineage tree.

#### Simulation 1

Our first example is a simple bifurcation of a single progenitor cell type into two descendant cell types (Fig. 4a). This is one of the simplest trajectory structures to recover and one where ordinary state-based inference already does well. Given a sufficiently accurate tree, LineageOT performs marginally better at ancestor prediction (Fig. 4b) and marginally worse at descendant prediction (Fig. 4c). In hindsight, this is not surprising. The lineage tree, rather than providing substantial new information, just reaffirms the natural assumption that cells in the same cluster are a bit more closely related.

#### Simulation 2

Inferring whether a single differentiated cell type came from multiple lineages is a common problem [27] and one of the standard goals of lineage tracing methods [32]. These convergent trajectories are difficult for state-based trajectory inference, which cannot distinguish the different ancestries of cells with similar measured states. Here we simulate two clusters that each split; after the split, two of the resulting clusters merge together (Fig. 4d). Now lineage information is important: LineageOT can separate cells in the convergent cluster by ancestry, while state-based methods cannot. Incorporating lineage information leads to substantially better prediction of ancestors than purely state-based optimal transport (Fig. 4e), without undermining descendant prediction (Fig. 4f).

#### Simulation 3

We included two unmerged clusters at the late time in Simulation 2 to illustrate how lineage information can resolve ambiguity: cells whose ancestry is unclear from expression alone should be coupled similarly to their close relatives in the lineage tree with unambiguous ancestry. The existence of separate unconverged clusters is not, however, necessary for separating cells by ancestry. In Simulation 3, we show two clusters that converge to a single final cell type (Fig. 4g). The distributions of descendants from each early cluster overlap too much for state-based methods like Waddington-OT to accurately infer ancestors. Despite the overlap, the descendant distributions remain sufficiently distinct for LineageOT to have nearly perfect assignment of late cells to early clusters, and thereby low ancestor error (Fig. 4h) with no loss of accuracy in descendant prediction (Fig. 4i).

#### Simulation 4

Our fourth example illustrates that lineage information can go beyond resolving ambiguity and even correct mistakes from state-based inference. We consider two clusters that split so that two of the late-time clusters end up closer to early cells that are not their ancestors (Fig. 4k). Optimal transport fails dramatically in this case, mapping entire clusters to the wrong set of ancestors. The failure is not due to any mistake in the algorithm: any method that uses only state information could not correctly infer the trajectory from this data. LineageOT, on the other hand, can use the shared ancestry to match clusters correctly, leading to significantly better prediction of both ancestors and descendants (Fig. 4l-m).

For this example, increasing the temporal resolution by sampling the system in between the two time points could allow optimal transport or other state-based methods to accurately describe the trajectories. In real biological systems with this type of ‘curled’ trajectory, lineage tracing may limit the need for many expensively-sampled timepoints, though it does add other costs like integrating the barcode editing technology into the genome. For convergent trajectories like Simulation 2 or 3, adding more time points without lineage information would not be enough: in that setting, lineage tracing is necessary for correct trajectory inference.

## Discussion

Analyzing the trajectories cells traverse during differentiation is crucial for understanding development and for harnessing the potential of stem cell therapies. However, general-purpose techniques for directly measuring trajectories of cellular differentiation remain elusive. In certain biological contexts, such as hematopoeisis [34] where cells are grown in suspension, daughter cells can be split and measured at different time-points. These direct connections between time-points allow for improved trajectory inference [18]. Such specialized techniques, however, are not applicable to systems where cells are adherent, because splitting daughter cells would perturb the developmental process. Trajectory inference from independent snapshots remains the most promising approach for understanding the genetic and epigenetic forces driving development in diverse biological contexts.

We develop the first general-purpose method for inferring developmental trajectories from scRNA-seq time-courses equipped with lineage information each time point. Lineage tracing techniques are progressing from early demonstrations of the technology [3, 26] through elaboration of the potential value of the data [32] and on towards future widespread use. We envision that scRNA-lineage time-courses will soon replace traditional scRNA-seq time-courses, because adding lineage information enables a far more powerful form of trajectory inference.

We demonstrate that LineageOT dramatically outperforms Waddington-OT on a time-course of *C. elegans* development (Fig. 3, Fig. S2, Fig. S3), and we illustrate through simulation that LineageOT can accurately recover complex trajectory structures that are impossible to recover from measurements of cell state alone (Fig. 4d-i). Effectively using the new lineage information experimentally accessible within each time point intuitively ought to improve inference accuracy across time points. LineageOT realizes that implicit potential.

Lineage trees are particularly helpful for untangling *convergent* trajectories, where cells arrive at a particular state from multiple ancestries. This occurs, for example, in the development of the lymphatic endothelium [27], macrophage development from embryonic or monocyte-derived progenitors [30], mouse gut endoderm [16], and several tissues in *C. elegans* ([17], Figs. 4a, S8, and S17). While finer temporal resolution might allow state-based trajectory inference to succeed in some of these examples, LineageOT can achieve higher accuracy with fewer time points. The couplings we recover enable a direct, rigorous approach to answer biological questions about ancestor-descendant relation in developmental processes and predict regulators that govern those transitions, as demonstrated by [24].

Our algorithm is derived from a flexible mathematical framework that can be adapted to include future methodological advances. Most immediately, novel methods for inferring a lineage tree from any kind of experiment, or from prior knowledge, can be used directly in the LineageOT pipeline. To leverage this to its fullest extent, one could incorporate an explicit quantification of uncertainty in the lineage tree. Furthermore, there could be significant advantages to simultaneously inferring the lineage tree together with the trajectories, rather than first fitting the tree and subsequently recovering a coupling. Finally, it might be possible to incorporate additional information, beyond cell state and cell lineage. For example, measurements of RNA velocity [10] could be incorporated into our framework of estimating ancestor or descendant states and then coupling across time points. As with LineageOT, the resulting algorithm would apply optimal transport with a modified cost function.

All of these improvements would build on the key observation that lineage tracing allows us to share information across closely related cells. State-based trajectory inference relies exclusively on the assumption that each descendant considered individually should be close in state to its ancestor. As we have demonstrated, expanding that assumption to consider related cells together allows for more powerful trajectory inference that can recover more complicated trajectories without relying on the restrictive assumption that cells with similar states having similar ancestry. LineageOT analyses of future cell state and lineage time-courses collected with current technologies will provide a new, more accurate window on the intricate processes of development.

## Acknowledgements

The authors are grateful to, among many others, Ruth Baker for helpful discussions and comments. AF is supported by the Royal Commission for the Exhibition of 1851. GS is supported by a Career Award at the Scientific Interface from the Burroughs Wellcome Fund, an NFRF Exploration Grant, and an NSERC Discovery Grant. We also thank the reviewers for their numerous helpful comments.

## Code availability

A demonstration implementation of LineageOT is available at https://github.com/aforr/LineageOT-demo.

## Data availability

*The C. elegans* data is available on GEO with accession code GSE126954 (https://www.ncbi.nlm.nih.gov/geo/query/acc.cgi?acc=GSE126954).

## Methods

### 1 State couplings and lineage couplings

A *developmental stochastic process* is a mathematical representation of a population of cells developing over time, where a single cell is represented by a point in a high-dimensional vector space of cellular states (e.g. gene expression space), and a population of cells is represented by a probability distribution on this state space. When we profile the population with scRNA-seq, we model the resulting data as a set of random samples from this probability distribution. In the context of development, a time-varying distribution ℙ_*t*_ represents the cells alive at time *t*, and the data from a scRNA-seq time-course consists of samples from ℙ_*t*_ collected at various times *t*_1_, *t*_2_, …, *t*_*N*_. The crucial point is that the random samples from different time points are *independent* in the probabilistic sense, because each time point is typically collected from a separate biological sample.

This brings us to the second key concept of a developmental stochastic process: the notion of a coupling connecting a pair of time points. We distinguish between two kinds of couplings: *state couplings* and *lineage couplings*. Intuitively, the state coupling connecting time *t*_1_ to *t*_2_ specifies relationships between ancestral states at *t*_1_ and descendant states at *t*_2_. Mathematically, it is a joint probability distribution over pairs of cell states (*x, y*), with *x* and *y* corresponding to cells alive at *t*_1_ and *t*_2_ respectively. Conditioning on cell state *x* at time *t*_1_ gives a distribution over possible descendant states *y* at time *t*_2_. In other words, while ℙ_*t*_ simply describes the states of cells that exist at each time point, the state couplings specify the trajectories that give rise to the changes we observe in the population. The state couplings contain information lost in a scRNA-seq time-course: the measurements are destructive so we cannot simultaneously measure the state of a cell and the state of its ancestors or descendants.

Even state couplings, however, still omit some of the information from a specific experiment or realization of the stochastic process. A cell *j* at time *t*_2_ has a true history, which may differ from the average history of cells with state equal to *y*_*j*_. Lineage information makes it possible to recover the history of *j* in particular. The history can again be described by a coupling, this time thought of as a coupling on cells rather than on states. We refer to this coupling as a *lineage coupling*.

One reason the distinction matters is that our descriptions of cell state are incomplete. Gene expression profiles, for example, are only one easily measured part of the cell state. Cells with similar current gene expression but different history could in principle differ in other aspects of their current state. Investigating that possibility requires separating the cells by ancestry even when their current states are similar.

### 2 Stochastic differential equation model

We consider a cell state at time *t* to be a point *x*(*t*) in some high-dimensional space 𝒳 such as gene expression space. Over time, cells follow some true path through 𝒳 according to a stochastic differential equation combining diffusion and drift:

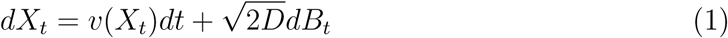

where *v* denotes a velocity field and *B*_*t*_ denotes standard Brownian motion scaled by the diffusion constant *D*. Note that diffusion dominates drift on short time-scales because diffusion is 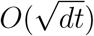 and drift is *O*(*dt*).

In this setting, we can model an experiment as sampling a set of cell paths {*x*_*i*_} from a distribution 𝒫 over the space of paths [0, 1] → 𝒳. Importantly, these paths are not observed in full; we only see *x*(*t*_1_) for the one measurement time *t*_1_. In a time-course experiment, in addition to measuring a set of cells {*x*_*i*_} at time *t*_1_ we also measure a second set of identically prepared cells {*y*_*j*_} at time *t*_2_.

We then want to couple the early and late distributions in order to trace cells forward and backward in time. As described above, a coupling *γ* is a joint distribution over pairs (*x*_*i*_, *y*_*j*_). When {*x*_*i*_} and {*y*_*j*_} are discrete sets, as they are here, *γ* is a matrix whose entries sum to 1.

The forward and backward questions are in principle different for lineage couplings. We could seek either a coupling *γ*^*F*^ such that 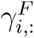, considered as a distribution on the {*y*_*j*_}, is approximately the true distribution of the descendants of cell *i*; or we could seek a coupling *γ*^*B*^ such that 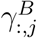, considered as a distribution on the {*x*_*i*_}, is approximately the true distribution of the ancestors of cell *j*. For one cell, that true ancestor distribution will be a single point mass.

### 3 Optimal transport as maximum likelihood estimate

For both the forwards and backwards problems, entropic optimal transport can be understood as the maximum likelihood coupling between an infinite population of cells started with the distribution of {*x*_*i*_} and conditioned to end up with the distribution of {*y*_*j*_}. If the likelihood of a cell at *x* ending at *y* is 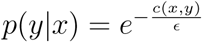, maximizing the log-likelihood log(*p*(*y*|*x, γ*)) leads to

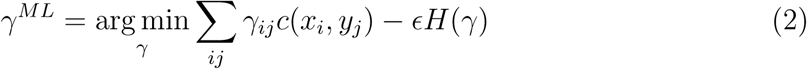

where *H*(*γ*) = − ∑_*ij*_ *γ*_*ij*_ log(*γ*_*ij*_) is the entropy of *γ*. This is precisely the objective function for optimal transport with cost *c*(*x, y*) and entropy parameter *ϵ*.

If the times *t*_1_ and *t*_2_ are sufficiently close together, the dynamics of *x* between *t*_1_ and *t*_2_ are approximately purely diffusive, so that *x*(*t*_2_)−*x*(*t*_1_) ∼ 𝒩 (0, *D*(*t*_2_ − *t*_1_)). This then translates to a quadratic optimal transport cost

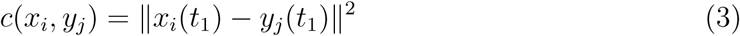

and entropy parameter

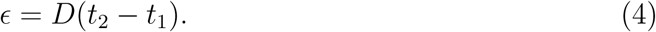

Because the likelihood is symmetric there is no difference between estimating forwards and estimating backwards. Other assumptions about the dynamics of the cells, such as might come from RNA velocity, could be incorporated here. Our goal with LineageOT is to find an appropriate replacement for the likelihood using the lineage information and use that as a new cost for optimal transport.

### 4 Ancestor inference with lineage information

A complete lineage tree 𝒯 for {*y*_*j*_} encodes the time 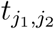 of the most recent common ancestor of each pair of cells 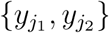. In terms of paths 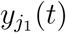 and 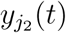 of the cells, a common ancestor at time 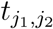 implies that

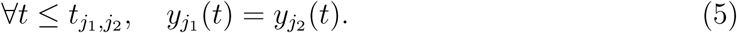

This gives no direct information about the unknown {*y*_*j*_(*t*_1_)}; instead, it tells us something about the correlations among {*y*_*j*_(*t*_1_)}. For LineageOT, we follow the same maximum-likelihood derivation that leads to entropic optimal transport but replace the distribution of *x* conditional on *y*_*j*_ with the distribution of *x* conditional on the full sample {*y*_*j*_} and the lineage tree 𝒯:

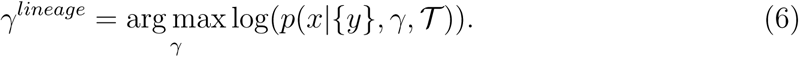

In the diffusive model where the differences in gene expression over time are Gaussian, if we are given the tree 𝒯 for {*y*_*j*_} then the expression values at the nodes (i.e., the common ancestors) are sampled from a Gaussian graphical model on the tree. We can then condition on the observed values {*y*_*j*_(*t*_2_)} to find the posterior density *p*(*y*_*j*_(*t*_1_)|{*y*_*k*_(*t*_2_)}, 𝒯). This density will be Gaussian with each mean 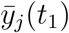 equal to a weighted average of the values {*y*_*j*_(*t*_2_)}. We then use an entropically regularized optimal transport coupling between 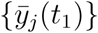 and {*x*_*i*_(*t*_1_)} to approximate the backwards coupling *γ*^*B*^.

Specifically, LineageOT implements the following procedure:

1. Fit a lineage tree estimate 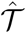 for {*y*_*j*_(*t*_2_)} including the estimated time of division of each most recent common ancestor, for example via neighbor-joining on CRISPR barcodes.
2. Add nodes {*y*_*j*_(*t*_1_)} for the ancestor of each time *t*_2_ cell at time *t*_1_ to 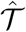. Some cells may share an ancestor here.
3. Pick a reference cell *y*_0_(*t*_2_). The difference in expression of other nodes of 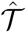 with respect to this reference (i.e., *y*_*v*_ − *y*_0_(*t*_2_)) is assumed to be normally distributed with mean zero; the precision matrix has entries

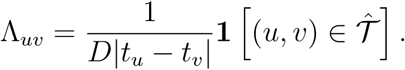
4. Condition on the values *y*_*j*_(*t*_2_) for the set 𝒪 of observed nodes. The conditional means for *y*_*v*_−*y*_0_(*t*_2_) in the set 𝓊 = 𝒪^*c*^ of unobserved nodes can then be found using the appropriately truncated precision matrix:

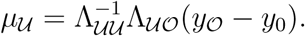
5. Compute the entropic optimal transport between {*x*_*i*_} and {*y*_*j*_} with cost

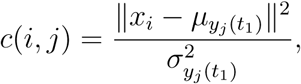

where 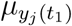and 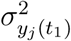 are the conditional mean and variance respectively for the ancestor of each *t*_2_ cell at time *t*_1_.

In practice, despite being designed for ancestor prediction rather than descendant prediction, LineageOT outperforms entropic optimal transport on both tasks for all but the simplest trajectories.

### 5 Fitting a lineage tree

To apply LineageOT, we need to infer a lineage tree that will define the structural equation model. We do not optimize this step, instead relying on a heuristic algorithm called neighbor joining [21]. Neighbor joining starts from pairwise lineage distance estimates, which can be estimated in CRISPR-based barcoding approaches using the Hamming distances between observed barcodes [3]. The fitted tree will not be perfect, and indeed simulations with currently plausible experimental parameters find significant errors in the inferred tree topology [23]. As our own simulations demonstrate, however, an imperfectly inferred tree can still substantially improve trajectory inference. Moreover, the source of the tree does not matter: a lineage tree based on detailed prior biological knowledge, as is available for *C. elegans*, can be used directly in LineageOT.

For LineageOT, we need not only the tree topology but also the time elapsed along each edge of the tree. The raw lineage distances computed from Hamming distances, however, give very noisy estimates of the edge times. We therefore correct the distances using the fact that all cells were sampled at the same time; this means that all leaves of the tree must have the same total distance to the root. Minimizing the mean squared error to the Hamming distance estimates subject to this constraint is a quadratic program that can be solved with standard convex optimization techniques and dramatically improves the estimated lineage distances (Fig. S7).

### 6 Error metrics

While we only produce one estimated coupling for both ancestor and descendant prediction, we separate out the two questions in evaluation. Given a true coupling *γ*^∗^, we define the *descendant prediction error* ℒ^*D*^(*γ*) for a fitted coupling *γ* with the same marginal over {*x*_*i*_} as the mean squared optimal transport distance between *γ*_*i*,:_ and 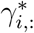 considered as distributions over {*y*_*j*_}:

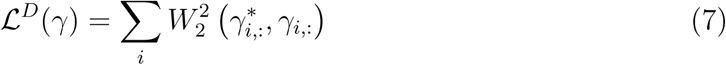

where *W*_2_(*µ, ν*) denotes the optimal transport distance between distributions *µ* and *ν* with quadratic cost, also called the Wasserstein-2 distance. Symmetrically, we define the *ancestor prediction error* ℒ^*A*^(*γ*) for a fitted coupling *γ* with the same marginal over {*y*_*j*_} as the mean squared optimal transport distance between *γ*_:,*j*_ and 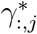 considered as distributions over {*x*_*i*_}:

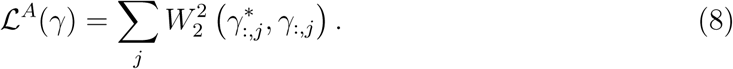

### 7 *C. elegans* ground truth and growth rates

Our ground truth coupling *γ*^∗^ for the *C. elegans* time-course is the forward coupling based on the lineage labels: we set 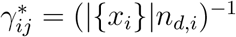, where *n*_*d,i*_ is the number of descendants of cell *x*_*i*_ in {*y*_*j*_}. This forward coupling has a uniform marginal over {*x*_*i*_} but not over {*y*_*j*_}. For simplicity, rather than using soft marginal constraints with estimated growth rates as

Waddington-OT does, we use the true marginals of *γ*^∗^ for all fitted couplings. Knowledge of the true marginals should help Waddington-OT and LineageOT approximately equally without significantly affecting the comparison between them.

### 8 Simulations

For our simulations, we construct a vector field to recreate a biologically plausible trajectory structure. Cells follow the vector field with diffusion and occasional cell division; the time between cell divisions is nearly constant, with normally distributed variability with a small variance (so that all sampled cell lifetimes are positive). Changing this variance in cell division times or setting it to 0 does not significantly affect our results. Meaningful dynamics occur in either two dimensions (for Simulations 1, 2, and 3) or three dimensions (for Simulation 4). The vector field is always constant in the first dimension, making *x*_1_ a proxy for time since the start of the simulation. In the remaining nontrivial one or two dimensions, we set the vector field to be the negative gradient of a potential with minima at locations we would like to have clusters, with the minima changing with *x*_1_. We simulate in three dimensions in all cases; for Simulations 1, 2, and 3, in the third dimension cells diffuse with no mean velocity. Thus, for example, the simple bifurcation of Simulation 1 follows the flow field

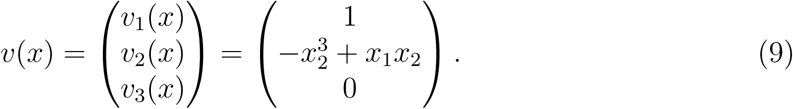

Initially, with *x*_1_ < 0, *x*_2_ = 0 is the only stable value of *x*_2_; later, with *x*_1_ > 0, there are two stable states 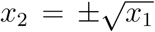. For the remaining simulations, which involve more complex piecewise-smooth flow fields, we refer readers to our code:

> https://github.com/aforr/LineageOT-demo.

Each cell has a lineage barcode that randomly mutates and is inherited by the cell’s descendants. The global mutation rate *r* is set so that the expected proportion of sites in a barcode of length *ℓ* that are unmutated at the time of sampling *t*_2_, equal to 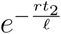, is close to 0.5 and so relatively far from both 0 and 1. The rate is then neither so slow that little lineage information is recorded so fast that barcodes are saturated before the sampling time. This choice was inspired by similar numbers in experimental data from [3].

For each vector field, we simulate a single embedded lineage tree measured at two time points and compute the couplings inferred by Waddington-OT, LineageOT given the true lineage tree, and LineageOT given a lineage tree fitted to the simulated barcodes. Because the simulated division rates are uniform across cells, we set the marginals for each fitted coupling to be uniform rather than inputting the true marginals as we did for the *C. elegans* evaluations. The fitted couplings are compared to the true coupling with the same ancestor and descendant prediction errors we used for *C. elegans*.

