## Supplemental Material for "A Unified Framework for Lineage Tracing and Trajectory Inference"

February 5, 2021

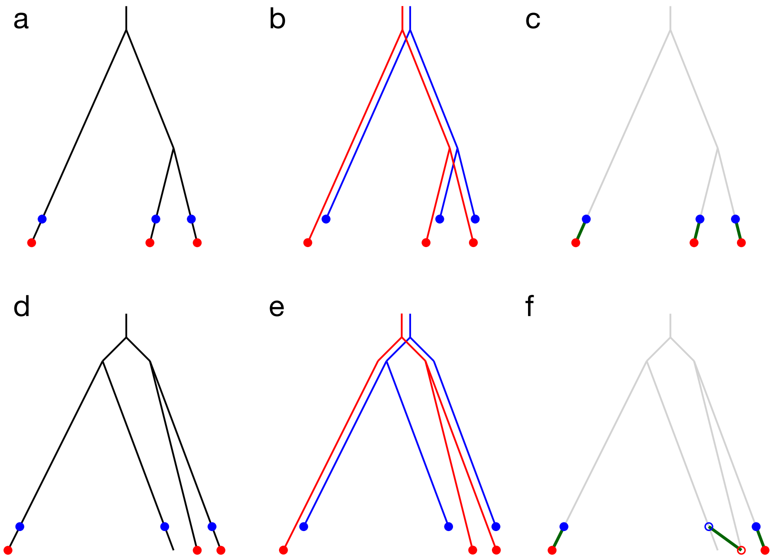


Figure S1: Due to subsampling or possible biological variability, patching together lineage trees from multiple time points is not always possible. (a) Ideally, measurements of cells at time $t_1$ (blue circles) and at time $t_2$ (red circles) would cover all branches of the same lineage tree (black). (b) In such a case, the lineage tree at $t_2$ (red) is an extended version of the lineage tree at $t_1$ (blue). (c) An ideal coupling of cells from $t_1$ to cells from $t_2$ then extends the lineage tree from $t_1$ to match the lineage tree at $t_2$. (d) Realistically, however, there may be some branches of the lineage tree measured at $t_1$ but not at $t_2$ and vice versa. (e) In such a case, the topologies of the lineage trees on the observed cells do not match up. The coupling that best matches cells at $t_1$ and $t_2$ to cells with similar expression to their descendants or ancestors respectively, shown in (f), does not patch the lineage trees together. A matching that preserved the topology of the observed lineage trees in this example would erroneously match the farthest left red cell to the farthest right blue cell.

### 1 Alternative evaluations of LineageOT on *C. elegans* data

Here we present comparisons of LineageOT and Waddington-OT for the two additional filtering strategies described in the main text. Computing the ancestor and descendant prediction errors is computationally expensive: given $n$ samples at each time point, $\mathcal{L}^D$ and $\mathcal{L}^A$ each require solving $n$ $n \times n$ optimal transport problems for a total cost of $O(n^3)$. The precisely annotated cells (Fig. 3, main text) and the ABpxp sublineage (Fig. S2) have few enough cells (151 and 2195 for the smallest and largest timepoints, respectively) that computing exact errors is still feasible. When we imputed precise lineages, however, we had up to 13000 cells per time point. To reduced computational effort, we computed couplings using all available cells and selected $k = 50$ cells at random from each true coupling marginal for the error calculations, reducing the computational cost to $O(kn^2)$. The results (Fig. S3a,c,d) are thus only approximate. The computational difficulty here is only for evaluation: applying LineageOT to new datasets without comprehensively comparing to a known ground truth requires much less effort.

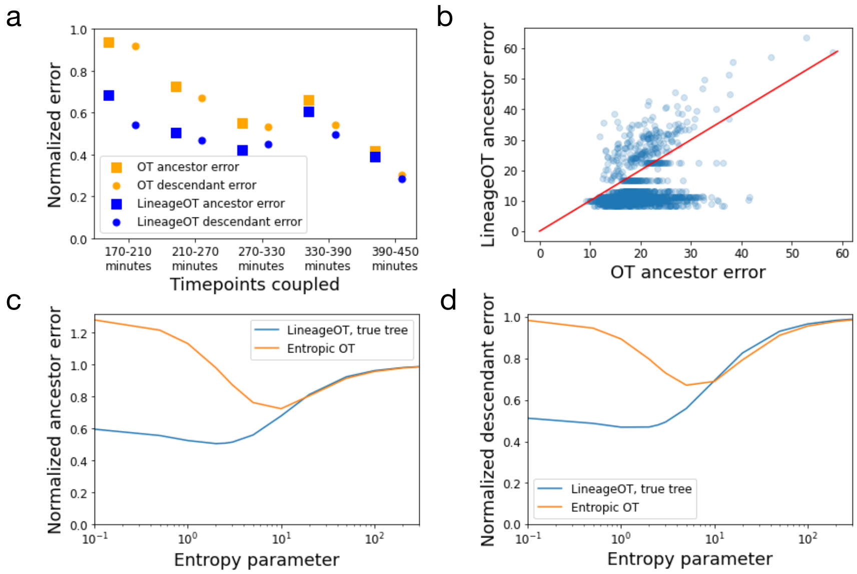


Figure S2: LineageOT outperforms optimal transport on the ABpxp sublineage. (a) Relative accuracy of optimal transport and LineageOT on all ABpxp cells and their descendants. Errors were normalized by dividing by the error of the noninformative independent coupling. (b) The error in predicting ancestor states, like the error for predicting descendant states (Fig. S4b), is lower for most cells with LineageOT. Here each point represents one cell from the 270 minute time point, which was coupled to the 210 minute time point. The red line marks equal error for both methods. LineageOT consistently improves on Waddington-OT for reasonable values of the entropy parameter, both in ancestor error (c) and descendant error (d), shown here for the 210-270 minute couplings. For each method in both (a) and (b), we chose the entropy parameters that gave the minimum error from parameter scans like those in (c) and (d).

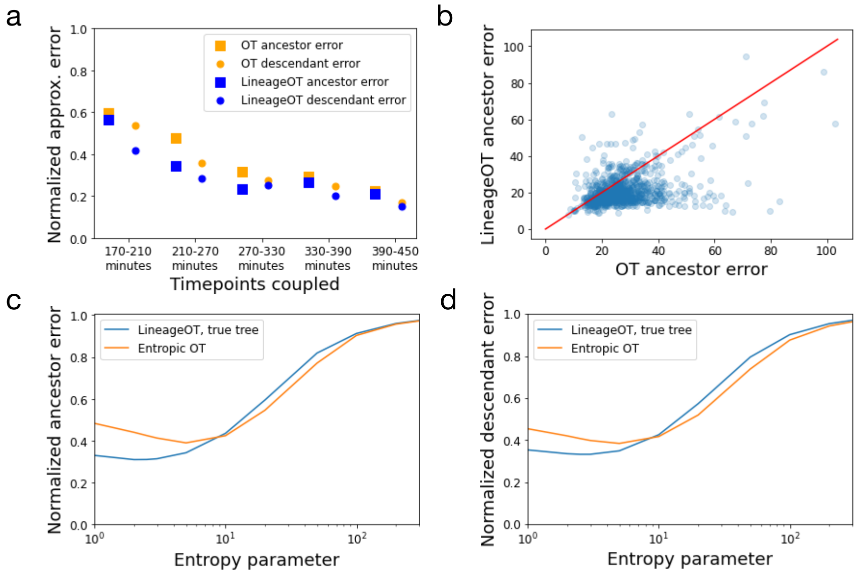


Figure S3: LineageOT outperforms optimal transport even with randomly imputed lineage annotations. (a) Relative accuracy of optimal transport and LineageOT on all cells with any lineage annotation. Errors were normalized by dividing by the error of the noninformative independent coupling. (b) The error in predicting ancestor states, like the error for predicting descendant states (Fig. S4c), is lower for most cells with LineageOT. Here each point represents one cell from the 270 minute time point, which was coupled to the 210 minute time point. The red line marks equal error for both methods. LineageOT consistently improves on Waddington-OT for reasonable values of the entropy parameter, both in ancestor error (c) and descendant error (d), shown here for the 210-270 minute couplings. For each method in both (a) and (b), we chose the entropy parameters that gave the minimum error from parameter scans like those in (c) and (d). In (a), (c), and (d), we compute average errors over a randomly selected subset of 50 cells, while in (b) we only plot 1000 randomly selected cells out of 8417 cells in the 270 minute batch.

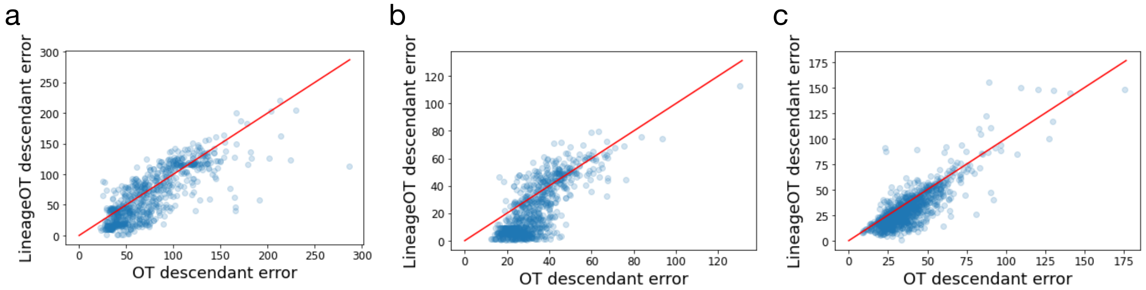


Figure S4: LineageOT tends to have lower descendant error for all three of our cell filtering methods. Here we compare errors for couplings between 210 and 270 minutes (a) restricting to precisely annotated cells, (b) restricting to the ABpxp subtree, and (c) randomly imputing precise lineage annotations. For (c) we only plot 1000 randomly selected cells out of 4147 cells in the 210 minute batch.

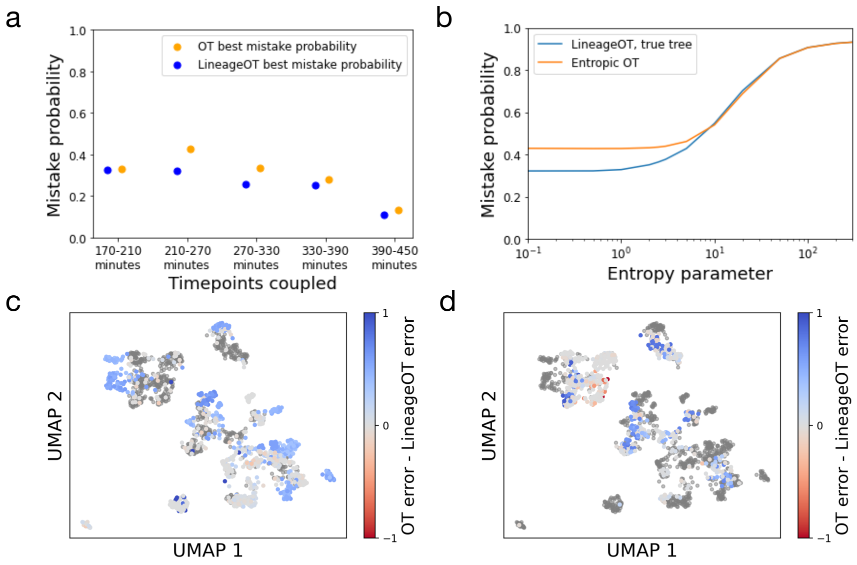


Figure S5: The improvement from LineageOT is robust to different choices of error metric. Here we measure performance as the probability of matching a cell to another cell that is not its ancestor or descendant, without using gene expression in evaluation. Mathematically, that is the total variation distance from an estimated coupling to the ground truth coupling after cells with identical lineage labels are identified. (a) Relative mistake probability of optimal transport and LineageOT on all cells with any lineage annotation. For this metric, the error for backwards prediction averaged over late cells is the same as the error for forwards prediction averaged over early cells. There is thus only one point per method and pair of timepoints. (b) As with the previous error metric, LineageOT matches or improves on Waddington-OT's performance for reasonable values of the entropy parameter, shown here for the 210-270 minute couplings. (c-d) In the same UMAP of 210 and 270 minute cells as Fig. 3 of the main text, late (c) or early (d) cells are colored by mistake probability from Waddington-OT minus the mistake probability from LineageOT. Blue indicates better performance by LineageOT, red better performance by Waddington-OT, and light grey approximately equal performance. The cells from 210 minutes and 270 minutes in (c) and (d) respectively are dark grey. For each method in both (a), (c), and (d), we chose the entropy parameters that gave the minimum error from parameter scans like the one in (b).

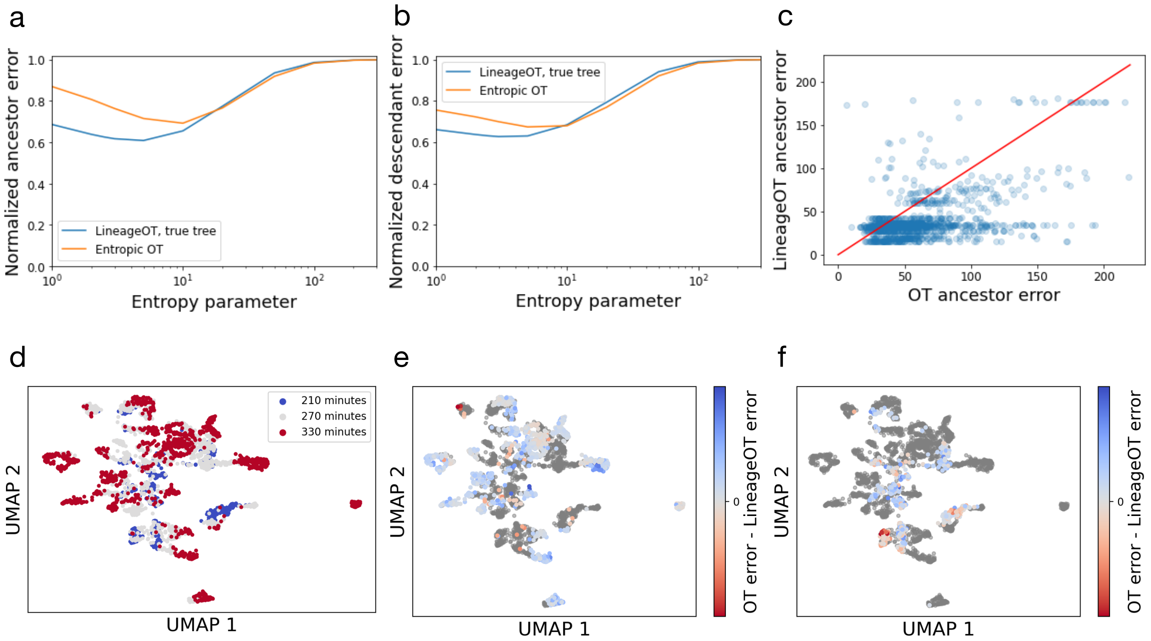


Figure S6: The improvement from adding lineage information persists when multiple couplings are concatenated. Here we show results from combining couplings of cells with complete lineage annotations from 210 minutes to 270 minutes and from 270 minutes to 330 minutes into couplings from 210 minutes to 330 minutes. Depending on the entropy parameter, LineageOT matches or exceeds the performance of Waddington-OT in both ancestor error (a) and descendant error (b). Errors were normalized by dividing by the error of the noninformative independent coupling. (c) The error in predicting ancestor states, like the error for predicting descendant states, is lower for most cells with LineageOT. Here each point represents one cell from the 330 minute time point, which was coupled to the 210 minute time point. The red line marks equal error for both methods. (d) UMAP visualization of the cells from the 210 (blue), 270 (grey), and 330 minute (red) time points. (e-f) Here, in the same UMAP, cells are colored by the ancestor (e) or descendant (f) error from Waddington-OT minus the same error from LineageOT. Blue indicates better performance by LineageOT, red better performance by Waddington-OT, and light grey approximately equal performance. The cells from 210 minutes and 330 minutes in (e) and (f) respectively, as well as the cells from 270 minutes in both plots, are dark grey as the corresponding error metric does not apply to them. For each method in (c), (e), and (f), we chose the entropy parameters that gave the minimum error from parameter scans like those in (a) and (b).

### 2 Tree fitting

A previous simulation study [1] showed that neighbor joining with currently-plausible experimental parameters recovered lineage trees from barcodes with only moderate accuracy according to the Robinson-Foulds distance. Robinson-Foulds, however, is a stringent error metric that penalizes small topological differences harshly. Perfect recovery according to Robinson-Foulds is not required for accurate downstream results. For LineageOT, the accuracy of pairwise lineage distances, defined for a pair of cells as twice the time to their most recent common ancestor, better characterizes the tree's accuracy. These lineage distances are closely related to the amount that information is shared across pairs of cells in LineageOT. By appropriately estimating edge lengths of the tree (Methods 5), we can get highly accurate lineage distance estimates even with relatively low Robinson-Foulds accuracy. We check the importance of tree accuracy in all our simulated examples by comparing the accuracy of LineageOT with a tree fitted to the barcodes and the experimentally unobserved true tree.

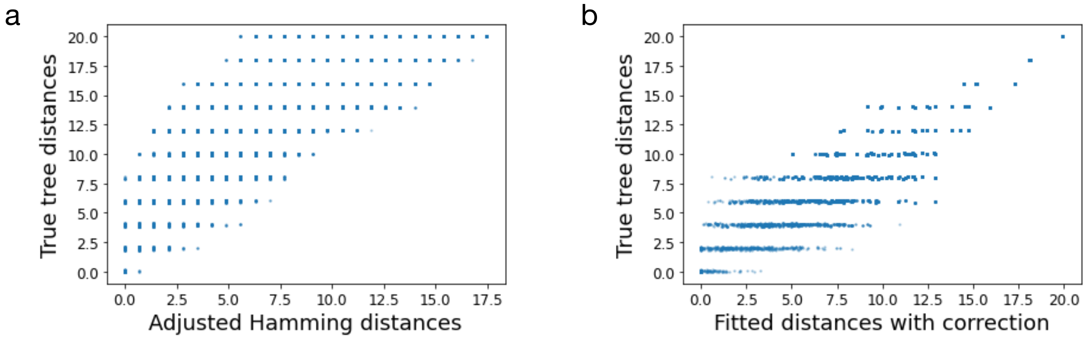


Figure S7: Accurately estimating pairwise lineage distances is possible even if the fitted tree has low Robinson-Foulds accuracy. (a) In this simulation, the scaled Hamming distance between barcodes has a correlation of only 0.81 with the true lineage distances. (b) Adjusting the distance estimates using the fitted tree and the constraint that cell sampling times are known increases the correlation with the true distances to 0.96. The Robinson-Foulds accuracy of this fitted tree is only 0.66.

### 3 Geodesics in distribution space

Optimal transport finds the shortest possible path between the initial and final distributions; mathematically, this means following the shortest geodesic according to the optimal transport metric. In Simulation 3, that is a mistake for Waddington-OT, but not because the true trajectory is far from a geodesic. Locally, the true trajectory does minimize the distance travelled by cells. Globally, however, the geodesic followed by the true trajectory is not the shortest path between its endpoints. Adding lineage information reduces or eliminates the error. We can see this cleanly by plotting the true and inferred couplings with a smaller number of cells (Fig. S8a-c).

Each coupling $\gamma$ implicitly defines a family of distributions interpolating between the distribution of $\{x_i\}$ and $\{y_j\}$. Given a time $t$ between $t_1$ and $t_2$ and a sample $(x_i, y_j)$ from $\gamma$, let

$$z_{ij}(t) = x_i \frac{t_2 - t}{t_2 - t_1} + y_j \frac{t - t_1}{t_2 - t_1}. \tag{1}$$

The distribution $Z_\gamma(t)$ of $\{z_{ij}(t)\}$ continuously changes from the distribution of $\{x_i\}$ at $t = t_1$ to the distribution of $\{y_j\}$ at $t = t_2$. If $\gamma$ accurately captures the true cell dynamics, the path these interpolants $Z_\gamma(t)$ follow in the space of distributions on $\mathcal{X}$ will approximately match the path of the unobserved true distributions. We sought to compare the interpolated distributions visually by embedding the paths in two dimensions.

To create a clean visualization, we simulated the double bifurcation from Fig. 4g-i with a low division rate so there were four cells at each time point. We computed interpolated distributions at 100 intermediate time points for each of the Waddington-OT, LineageOT, and ground-truth couplings and then found the optimal transport distances between all pairs of interpolated distributions. Using that distance matrix, we embed the distributions in two dimensions using multidimensional scaling, an algorithm that attempts to place points in Euclidean space so that pairwise distances are preserved (Fig. S8d). Because the space of distributions with the optimal transport metric is both high-dimensional and non-Euclidean, the embedding necessarily has some distortion. Still, some features of the high-dimensional paths remain: notably, both paths are approximately straight lines near the start point, as expected for geodesics.

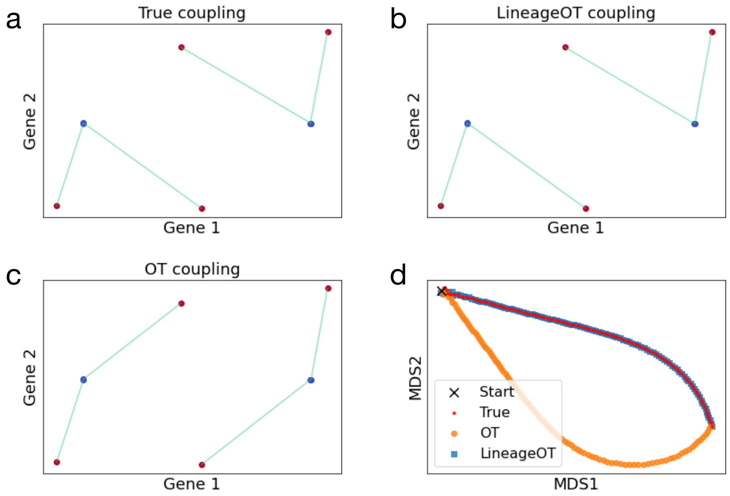


Figure S8: When the assumption that descendant states are closest to their ancestors is violated, Waddington-OT makes clear mistakes that can be corrected with lineage information. Here the cell dynamics are the same as in Fig. 4k except for a lower division rate which leads to a smaller number of cells. The true coupling (a) matches the LineageOT coupling (b) exactly, while Waddington-OT (c) mismatches the cells in the center. Early cells are shown in blue and late cells in red. (d) Visualization of distributions interpolated using the true (red), Waddington-OT (orange), and LineageOT (red) couplings. Each point is a distribution in between times $t_1$ and $t_2$ embedded in two dimensions using multidimensional scaling; the black $\times$ marks the initial distribution at time $t_1$, corresponding to the blue cells in (a-c). The interpolants for each coupling would follow a straight line if visualized individually; the curvature is a distortion required for a picture in two dimensional Euclidean space.

### References

[1] Irepan Salvador-Martínez, Marco Grillo, Michalis Averof, and Maximilian J Telford. Is it possible to reconstruct an accurate cell lineage using CRISPR recorders? *eLife*, 8, 2019.
